# Plasticity of the glutamate transporter EAAT2 on striatal astrocytes regulates flexibility in behavior

**DOI:** 10.1101/2020.01.30.927053

**Authors:** Arjen J. Boender, Raffaella Tonini

## Abstract

Modulation of striatal circuits is necessary for behavioral flexibility and confers the ability to adapt to environmental changes. Striatal astrocytes contribute to circuit neuromodulation by controlling the activity of ambient neurotransmitters. In particular, extracellular glutamate levels are tightly controlled by the astrocytic glutamate transporter EAAT2, thereby influencing synaptic functioning and neural network activity. While disturbances in EAAT2 have been related to neurological dysfunctions, it remains unclear if environmental cues influence this protein’s function to specifically shape action control.

In this study, we investigate the relationship between experience-dependent plasticity of EAAT2 expression and action inflexibility that follows overtraining for an instrumental task. We find that task overtraining is associated with the upregulation of EAAT2 in the lateral part of the dorsal striatum (DLS). Interfering with EAAT2 upregulation by chemogenetic activation of astrocytic Gq signaling or by transient *in vivo* knockdown of EAAT2 in the DLS restores behavioral flexibility. Astrocytes are emerging as critical regulators of striatal functions, and by demonstrating that plasticity of EAAT2 expression in the DLS shapes behavior, this work provides novel mechanistic insights into how flexibility in action control is regulated.

## Introduction

Astrocytes are the major glial cell type in the brain. They shape synaptic functions, establishing a tight functional interplay with neurons through the release of neuroactive substances (i.e. gliotransmitters) and regulating glutamate clearance (Araque et al., 2014; Bergles and Jahr, 1997; Clark and Barbour, 1997; Clements et al., 1992; Murphy-Royal et al., 2017; Perea et al., 2009). Recent studies indicate that astrocyte-neuron communication via gliotransmission occurs at specific synapses and drives defined behaviors (Martin-Fernandez et al., 2017; Sardinha et al., 2017). In contrast, the link between the regulation of glutamate clearance and behavioral outcomes is underexplored (Valtcheva and Venance, 2019). Astrocytes control glutamate clearance through the expression of excitatory amino acid transporters (EAATs) (Divito and Underhill, 2014). In particular, the glutamate transporter EAAT2 accounts for ~95% of the glutamate uptake capacity of the brain and is predominantly expressed in astrocytes (Danbolt, 2001; Petr et al., 2015). Deleting EAAT2 from astrocytes leads to epilepsy and self-mutilation, while neuronal EAAT2 deficiency has no severe behavioral consequences (Aida et al., 2015; Petr et al., 2015). These studies suggest a critical role for astrocytic EAAT2 in regulating behavior.

Astrocytic EAAT2 is critical for setting the strength of synaptic inputs and preserving the timing of coordinated presynaptic and postsynaptic neuronal activity (Min and Nevian, 2012; Valtcheva and Venance, 2016). These functions may be particularly relevant for procedural learning processes that precisely integrate sensory and motor information, such as operant conditioning. This behavior relies on two learning procedures and is supported by distinct anatomical substrates. During the initial acquisition, instrumental performance is goal-directed and depends on learning the causal relationship between actions (A) and their outcomes (O) through information processing at associative cortical inputs to the dorsomedial striatum (DMS). After repeated practice, behavior become inflexible, primarily through strengthening of stimulus-response associations (Balleine and Dickinson, 1998; Nazzaro et al., 2012; O’Hare et al., 2016; Peak et al., 2018; Shiflett and Balleine, 2011b; Yin et al., 2004, 2005a). Strengthening of these associations requires plasticity at projections from sensorimotor cortices to the dorsolateral striatum (DLS) (Gremel and Costa, 2013; Nazzaro et al., 2012; O’Hare et al., 2016; Thorn et al., 2010).

EAAT2 expression changes upon sensory experience (Genoud et al., 2006) and pharmacological overexpression of striatal EAAT2 affects time-coded plasticity at corticostriatal synapses (Valtcheva and Venance, 2016). However, it has not been explored whether striatal EAAT2 and its regulation during repeated instrumental conditioning influences action control. In this study, we specifically investigated the relationship between experience-dependent plasticity of EAAT2 levels in the dorsal striatum and the loss of behavioral flexibility induced by overtraining on an appetitive learning task (Nazzaro et al., 2012). We find that the inability to encode new A-O associations is associated with the upregulation of EAAT2 in the lateral part of the dorsal striatum (DLS), but not in the dorsomedial striatum (DMS). These behavioral and molecular effects were prevented by interfering with EAAT2 upregulation through the chemogenetic activation of astrocytic Gq signaling or by *in vivo* transient knockdown of EAAT2 in the DLS. This study thus demonstrates that astrocytes contribute to overtraining-induced behavioral inflexibility via regulating EAAT2 expression in the DLS.

## Material and Methods

### Experimental design and statistical analyses

All animal experiments were designed in accordance with the ARRIVE (Animal Research: Reporting of In Vivo Experiments) guidelines (Kilkenny et al., 2010), with a commitment to refinement, reduction, and replacement, and using power analyses to optimize sample size, as in our previous peer-reviewed work (Nazzaro et al., 2012; Trusel et al., 2015). Appropriate parametric statistics were used to test hypotheses, unless data did not meet the assumptions of the intended parametric test (normality test). In this case, appropriate non-parametric tests were used. Power analysis specifications to estimate sample size were: power = 0.8, alpha = 0.05, two-tailed, and an effect size that is 50% greater than previously observed standard deviations. Data were analyzed by two-way repeated measure ANOVA (RM2WA) or one-way repeated measure ANOVA (RM1WA) for comparisons within a group, and one-way ANOVA (1WA) for between-group comparisons (GraphPad Prism 6 software). A mixed-effects two-way ANOVA was used to analyze experiments with between-subjects (short- and overtraining) and within-subjects variables (post-training positive A-O versus negative A-O or valued versus devalued conditions) (IBM SPSS Statistics v.25 software). Corrected post-hoc tests (Tukey or Sidak as indicated) were performed only when the ANOVA yielded a significant main or interaction effect. Two groups were tested for statistical significance using the independent samples t-test, the paired samples t-test, or equivalent non-parametric tests (GraphPad Prism 6 software). Statistical details of experiments are shown in the results, figures and figure legends. Data are reported as mean ± s.e.m. Blinding was applied to treatment administration and data analysis.

### Animals and tissue processing

Male C57BL/6J mice at postnatal day 35-50 (Charles-River, Italy) were used for all experimental procedures and were housed 2-5 per cage in a temperature- and humidity-controlled room under a 12:12 h light/dark cycle with lights on at 07:00. After experiments, animals were deeply anesthetized followed by decapitation or transcardial perfusion with 0.1 M phosphate-buffered saline (PBS, pH 7.4) containing 4% paraformaldehyde (PFA). For subsequent qPCR and western blot analyses, brains were sectioned using a brain matrix (Zivic Instruments, IL, USA). Next, sections were rapidly frozen and the dorsolateral (DLS) and dorsomedial striatum (DMS) were dissected. For subsequent immunohistochemical analyses, PFA-fixed 40-μm brain sections were made on a cryostat (Leica, Germany). All experimental procedures were carried out according to the directives of the Italian Ministry of Health (D.Igs 26/2014) and the European Community (86/609/EEC) that regulate animal research.

### Viral infusion

Mice were anesthetized with a 1.5-4% isoflurane/O_2_ mixture and fixed in a stereotactic frame (Stoelting Inc., IL, USA). 150 nl of 1.0×10^9^ genomic copies/μl of AAV5-GFAP-HA-hM3Dq or AAV5-GFAP-eGFP (UNC Vector Core, NC, USA) were infused in the DLS (relative to Bregma: + 0.5 mm anteroposterior, ± 2.60 mm mediolateral and − 3.5 mm dorsoventral) at a rate of 25 nl/min. In each animal, viral infusion sites were determined on coronal slices that covered the full anterior-posterior range. The extent of DLS viral transduction was determined by calculating the average percentage of regional infection across all animals and subdividing the average values in groups of low (<35% of animals), moderate (35-75% animals) and high (>75% of animals) infected areas. Infection was observed in the DMS of one animal, which was therefore excluded from further analyses.

### Cannula placement and local infusions

Mice were anesthetized with a mix of 1.5-4% isofluorane/O_2_ mixture and fixed in a stereotactic frame (Kopf Instruments, Germany). Two 26-gauge stainless steel cannulas (UNIMED, Switzerland) were implanted and cemented to the skull at the following coordinates (relative to Bregma): + 0.5 mm anteroposterior, ± 2.60 mm mediolateral and − 2.5 mm dorsoventral. Dummy injectors (UNIMED) were placed to ensure patency. For local infusions, we used injectors (UNIMED) that protruded 0.1 mm from the guide cannula to target the infusions to the DLS. 300 nl of 10 μM MO solution (3 pmol; Gene Tools, LLC, OR, USA) was infused at a rate of 100 nl/min. Antisense sequences: Control MO, 5’-CCTCTTACCTCAGTTACAATTTATA-3’; EAAT2 MO, 5’-GCACCCTCTGTTGATGCCATG-3’. Cannula placement was determined on coronal slices that covered the full anterior-posterior range. The lowest dorsoventral point was taken as a proxy for injection location. In none of the animals were injection points located in the DMS. Animals that lost their cannulas over the course of the experiment were excluded from analyses (n=4).

### Behavioral experiments

Behavioral experiments took place in operant chambers (17.8 cm × 15.2 cm × 18.4 cm; Med Associates, VT, USA) equipped with two holes on either side of the food magazine, in which the food pellets were delivered. Mice were trained to nose poke in one of two holes to obtain food pellets (reinforcers), which could be either chocolate (F0 5301, Bilaney, UK) or sucrose (F0 5684, Bilaney, UK). One type of reinforcer was delivered in the operant chamber, contingent upon nose poking, while the other was presented in matched amounts non-contingently in the home cage. Magazine entries were recorded using an infrared beam. The type of reinforcer and the side of the active nose poke hole (only poking in the active nose poke led to pellet delivery) were counterbalanced. Before training, mice were food-deprived to maintain 85-90% of their initial body weight. Naive, short-trained and overtrained mice were food-restricted for the same amount of time, in order to control for putative effects of food restriction on EAAT2 expression. Mice were fed daily, directly after the training sessions with standard lab chow (Labdiet 5001, Labsupply, TX, USA) and one of two reinforcers (non-contingently). Initial nose poke training consisted of two daily sessions of continuous reinforcement (CRF), in which every active nose poke was rewarded until 10 reinforcers were earned. After the CRF sessions, mice were trained on variable interval (VI) schedules (Hilario et al., 2007; Nazzaro et al., 2012; Rossi and Yin, 2012), in which active nose pokes were reinforced after variable time intervals that lasted on average 30 s (VI-30) or 60 s (VI-60), and ended after 20 reinforcers. After three daily VI-30 sessions, mice were either short-trained for four daily VI-60 sessions or overtrained for 18 VI-60 sessions.

#### Post-training omission procedure

The omission test started one day after instrumental training and lasted two days. On day 1, mice were exposed to a 30 min control session under the prevailing, positive action-outcome contingency (nose poking leads to reinforcer delivery). On day 2, mice were subjected to a 30 min session in which the previously learned A-O contingency was reversed (negative contingency) in that the pellet was delivered every 20 s without nose poke, but each nose poke would reset the counter and delay the food delivery. The rates of active nose poke (ANP) under the two different A-O contingencies (negative A-O / positive A-O) were used to determine behavioral flexibility (Nazzaro et al., 2012; Rossi and Yin, 2012; Yu et al., 2009).

#### Post-training devaluation procedure

In the devaluation procedure, the outcome of nose poking (reinforcer delivery) was devalued using sensory-specific satiety (Hilario et al., 2007; Nazzaro et al., 2012). The devaluation test started 24 h after the last training session and lasted 2 days. On each day, mice were exposed *ad libitum* to one of the reinforcers (chocolate or sucrose) for 60 min in a separate cage. On day 1, mice were given the reinforcer, previously earned by nose poking (devalued condition); on day 2 mice received the reinforcers, previously available in their home cages during training (valued condition). The order of the valued and devalued conditions was randomized. Immediately after each feeding session, the mouse underwent a 5 min extinction test in the operant chamber, during which no reinforcer was delivered. The number of nose pokes into the active hole under the valued and devalued conditions were compared.

### Drugs

Water-soluble clozapine-N-oxide dihydrochloride (Tocris, United Kingdom), was diluted in sterile saline (0.9% NaCl) and administered i.p. (3 mg/kg). Vehicle (0.9% NaCl) was administered in equal amounts.

### Quantitative PCR

Total RNA was isolated with QiaZOL according to the manufacturer’s instructions (Qiagen, Germany). After DNAse treatment (Promega, WI, USA), complementary DNA was synthesized (cycle conditions: 25°C for 5 min, 42°C for 1 h, 70°C for 10 min) on a Peltier Thermal Cycler (Bio-Rad Laboratories Inc., Hercules, CA, USA) using the GoTaq^®^ 2-step RT-qPCR kit (Promega). Quantitative PCR runs were performed on a 7900HT Fast Real-Time PCR System (Applied Biosystems Inc., CA, USA) using the primer sets listed in Table 1 (cycle conditions: 2 min of initial denaturation at 95°C, followed by 40 cycles of 15 s at 95°C and 45 s at 60°C). We normalized expression values relative to *Gadph* and *Ppia* using BioGazelle (BioGazelle, Belgium).

### Western blot

For total protein extraction, dissected pieces of frozen dorsal striatal tissue were homogenized in ice-cold TEVP-buffer (pH 7.4, 10 mM Tris-HCl, 5 mM NaF, 1 mM Na_3_VO_4_, 1 mM EDTA, 1 mM EGTA and 320 mM sucrose). After sonication, we determined protein concentration using the BCA method (ThermoFisher Scientific, MA, USA), loading equal quantities (15 μg) per lane. Western blotting was performed with a 1:5000 concentration of primary rabbit anti-EAAT2 antibody (SAB2104141, Sigma-Aldrich). We normalized the expression of EAAT2 to the expression of primary rabbit anti-calnexin (ABI-SPA-860, Enzo Life Sciences Inc., NY, USA, 1:10000). Blots were incubated with horseradish peroxidase-conjugated anti-rabbit secondary antibody (1:10000; Amersham Biosciences, NJ, USA). Bands were detected using enhanced chemiluminescence (Pierce Biotechnology, IL, USA) and imaged with a LAS 400 Mini Imaging System (GE Healthcare, IL, USA). Immuno-positive bands were quantified using ImageJ software (NIH, MA, USA) and densitometric values of EAAT2 protein were normalized to calnexin protein values.

### Immunohistochemistry

Sections were blocked and permeabilized for 1 h in PBS containing 5% normal donkey serum and 0.05% Tween-20. After being washed in PBS, sections were incubated overnight with primary antibodies in PBS containing 0.05% Tween-20. ALDH1L1 was detected with rabbit anti-ALDH1L1 (1:500, AB87117, Abcam), cFOS was detected with guinea pig anti-cFOS (1:1000, 226004, Synaptic Systems GmbH, Germany), GFAP was detected with rabbit anti-GFAP (1:1000, Z0334, Agilent, CA, USA), HA-tagged DREADD receptors were detected with goat anti-HA (1:1000, AB9134, Abcam, UK), MAP2 was detected with rabbit anti-MAP2 (1:1000, AB32454, Abcam) and NeuN was detected with rabbit anti-NeuN (1:1000, AB104225, Abcam). After being washed again in PBS, sections were incubated with Alexa-labeled secondary antibodies for 1 h (1:500, Molecular Probes, USA). Sections were then washed in PBS and mounted in Fluoromount containing DAPI (Calbiochem, USA).

### Image analyses

Immunofluorescent sections were digitally imaged using an Olympus epifluorescent microscope (Olympus, Japan) equipped with the Neurolucida imaging system, a Keyence BZ-X800 microscope (Keyence, Japan) or a Nikon A1 confocal microscope (Nikon, Japan). To validate astrocytic expression of hM3Dq, we analyzed six animals (at least two slices per animal), and to examine CNO-induced cFOS expression, five animals. We determined the percentage of colocalization of ALDH1L1 with hM3Dq or MAP2 with hM3Dq using ImageJ (NIH; JaCOP plugin to determine colocalization) and the percentage of cFOS colocalization with hM3Dq+ cells or NeuN+ cells by manual counting. Injection sites were verified using four coronal slices that covered the full anterior-posterior range of the dorsal striatum. GFAP density after CNO and MO treatment was determined via the intensity of GFAP-immunofluorescence, normalized by area. The percentage of NeuN+ nuclei after MO treatment was determined by manual counting.

## Results

### Overtraining of instrumental conditioning induces inflexible behavior and upregulates EAAT2 in the DLS

To study experience-dependent plasticity of EAAT2 expression, we subjected mice to an instrumental task. Mice were either short-trained (Short) or overtrained (Over) to nose poke for food reinforcers under a variable interval schedule of reinforcement (Nazzaro et al., 2012; Rossi and Yin, 2012) (**Figure 1A**). All mice performed indistinguishably in the acquisition phase of instrumental learning: the rates between groups across sessions did not differ for active nose pokes (ANP) (Short n = 15, Over n = 14; ANP/min, session: F_8,216_ = 82.04, p < 0.0001; group: F_1,27_ = 0.02, p = 0.88; session X group interaction: F_8,216_ = 0.81, p = 0.6; **Figure 1B**), inactive nose pokes (INP), or magazine entries (ME) (p ˃ 0.05; **Figure S1A**).

**Figure 1.**
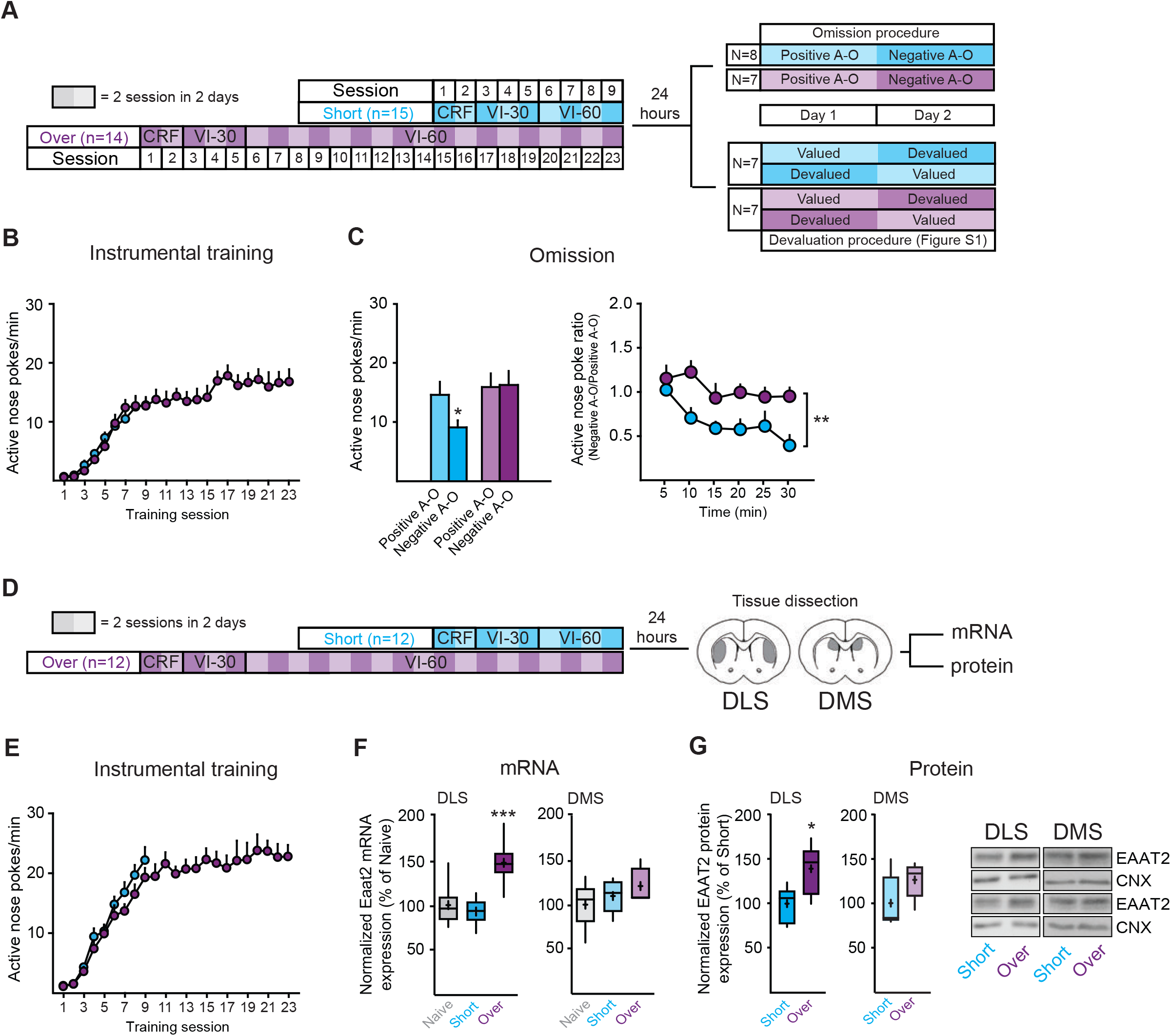
Repeated instrumental training leads to behavioral inflexibility and increases EAAT2 expression in the dorsolateral striatum (DLS). (**A**) Schematic illustrating the behavioral regimens (CRF: continuous reinforcement, VI-30: 30 s of variable intervals, VI-60: 60 s of variable intervals). (**B**) Active nose poke (ANP) rates during training in short- (n = 15) and overtrained (n = 14) mice. (**C**) Post-training omission procedure. (Left) Comparison of ANP rates under positive and negative contingency in both short- and overtrained mice (short-trained n = 8, positive contingency: 14.7 ± 2.2, negative contingency: 9.2 ± 1.2, Sidak: * p = 0.02; overtrained n=7, positive contingency: 16 ± 2.3, negative contingency: 16.4 ± 2.4; Sidak: p = 0.98). (Right) Time courses of ANP ratios (ANP rates under negative contingency/ANP rates under positive contingency) in short- and overtrained mice. (**D**) Schematic depicts the behavioral regimens followed by mRNA and protein analysis. (**E**) ANP rate increased during training in both short- and over-trained mice and did not differ between groups (short-trained n = 12, overtrained n = 12; session: F_8,176_ = 89.42, p < 0.0001; group: F_1,22_ = 2.07, p = 0.16; session X group interaction: F_8,176_ = 0.77, p = 0.63). (**F**) Box-whisker plots of normalized expression levels of *Eaat2* mRNA in the dorsolateral striatum (DLS) (Tukey, naive versus short-trained, p > 0.05; naive versus overtrained, ** p < 0.01; short-versus overtrained, ** p < 0.01) and dorsomedial striatum (DMS) of naive (cage control), short-trained and overtrained mice. In this and similar plots throughout the manuscript, the horizontal line indicates the median; the cross equals the mean; the box represents the 25^th^ and 75^th^ percentiles; and whiskers are the minimal and maximal values. (**G**) Box-whisker plot of normalized levels of EAAT2 protein in the DLS and DMS of short- and overtrained mice. Blots are representative samples from each group. CNX, calnexin. In the averaged time courses (**B**, **C**, **E**) and bar graphs (**C**), values are presented as mean ± S.E.

Performance in short-trained mice is expected to be controlled by flexible, goal-directed behavior, which is sensitive to changes in the prevailing action-outcome (A-O) contingency learned during training. In contrast, overtrained mice should show attenuated behavioral sensitivity to manipulation of A-O associations (Hilario et al., 2007; Nazzaro et al., 2012; O’Hare et al., 2016; Peak et al., 2018). To confirm these behavioral characteristics in our experimental setting, we measured behavioral sensitivity to changes in the prevailing A-O contingency by exposing a subset of mice to a post-training omission procedure in which the previously learned A-O association is reversed. During this procedure, pellets are delivered when mice refrain from nose poking and omitted when they nose poke (**Figure 1A** and **Figure 1C**) (Derusso et al., 2010; Nazzaro et al., 2012; Rossi and Yin, 2012; Yu et al., 2009). Nose poking behavior in short-trained mice was sensitive to changes in A-O contingency, shown by the reduced ANP rates over the post-training A-O reversal session (negative contingency) compared to the post-training control session (positive contingency; **Figure 1C**). This reduction in ANP did not occur in overtrained mice, which were insensitive to contingency reversal, suggesting inflexible behavior (Short n = 8, Over n = 7; ANP/min, post-training omission procedure: F_1,13_ = 3.93, p = 0.07; group: F_1,13_ = 2.71, p = 0.12; contingency X group interaction: F_1,13_ = 5.1, p = 0.04; Sidak, positive versus negative contingency: Short, * p = 0.02; Over, p = 0.98; **Figure 1C**). Accordingly, time course analysis of ANP ratios (ANP rate under negative contingency/ANP rate under positive contingency) revealed a main group effect (ANP ratio, time: F_5,65_ = 5.98, p = 0.0001; group: F_1,13_ = 9.89, ** p = 0.008; time X group interaction: F_5,65_ = 1.5, p = 0.2; **Figure 1C**). Moreover, while short- and overtrained mice earned a similar number of reinforcers under the positive A-O contingency, short-trained mice received more reinforcers than overtrained mice after A-O contingency reversal (obtained reinforcers, contingency: F_1,13_ = 22.17, p = 0.0004; group: F_1,13_ = 7.43, p = 0.017; contingency X group interaction: F_1,13_ = 6.95, p = 0.02; Sidak, short- versus overtrained: positive contingency, p > 0.05; negative contingency, ** p < 0.01; **Figure S1B**). Thus, the ability to adapt performance during the A-O reversal enabled short-trained mice to affect the task outcome (i.e. amount of obtained reinforcers) more efficiently than the inflexible behavioral strategy used by overtrained mice. The difference in behavioral flexibility between short- and overtrained mice cannot be attributed to a general suppression of behavior in short-trained animals, as the INP and ME rates did not differ between the two groups either before or after the reversal in A-O contingency (p ˃ 0.05; **Figure S1B**).

Behavioral inflexibility that follows task overtraining points to stimulus-driven habitual performance, which not only renders behavior impervious to changes in A-O contingencies (as shown), but also confers behavioral insensitivity to action-outcome devaluation (Balleine and O’Doherty, 2010; Malvaez et al., 2018; Nazzaro et al., 2012; Rossi and Yin, 2012). To further assess if the behavior of overtrained mice in our experimental setting was consistent with habitual performance, a second subset of short- and overtrained mice (**Figure 1A-B** and **Figure S1A, C**) was subjected to a devaluation test, in which the value of the action outcome (reinforcer delivery) was decreased by specific sensory satiety (Hilario et al., 2007; Nazzaro et al., 2012; Shiflett et al., 2010). As expected when behavioral performance is dominated by a flexible, goal-directed strategy, the rates of ANP of short-trained mice in the devalued condition decreased relative to the valued condition. In contrast, overtrained mice did not show any effect of devaluation (Short n = 7, Over n = 7; ANP/min; condition: F_1,12_ = 4.59, p = 0.05; group: F_1,12_ = 0.49, p = 0.5; condition X group interaction: F_1,12_ = 9.94, p = 0.008; Sidak, valued versus devalued condition: short-trained, ** p = 0.006; overtrained, p = 0.74; **Figure S1C**). Pellet consumption and INP and ME rates were similar in the valued and devalued conditions for both short- and overtrained mice (p ˃ 0.05; **Figure S1C**). These results are consistent with our previous observations of mice exposed to analogous training regimes (Nazzaro et al., 2012). In summary, our experimental conditions result in short- and overtrained mice that use distinct flexible and inflexible behavioral strategies, respectively.

We then assessed whether overtraining affects EAAT2 expression by measuring *Eaat2* mRNA and EAAT2 protein levels in the DLS and DMS of a new cohort of short-trained and overtrained mice, 24 h after their last training session (**Figure 1E-G** and **Figure S1D**). Compared to naive (home-cage controls) and short-trained mice, overtrained animals had higher *Eaat2* mRNA levels in the DLS (Naive, n = 8, 100% ± 7.8; Short n = 6, 93% ± 6.2; Over n = 6, 147.2% ± 10.6; *Eaat2* expression, 1WA, F_2,17_ = 11.51, *** p = 0.0007; **Figure 1F**), but not in the DMS (Naive n = 8, 100% ± 8.5; Short n = 6, 109.7% ± 7.6; Over n = 5, 120.9% ± 9; *Eaat2* expression, 1WA, F_2,16_ = 1.46, p = 0.26; **Figure 1F**). Similarly, EAAT2 protein levels were significantly higher in the DLS of overtrained mice than in short-trained mice (Short n = 6, 100% ± 8.1; Over n = 6, 139% ± 11.3; EAAT2 expression, Mann-Whitney test, U = 5, * p = 0.04, **Figure 1G**), whereas protein levels in the DMS were comparable between the two groups (Short, 100% ± 12; Over, 126% ± 8.3; EAAT2 expression, Mann-Whitney test, U = 8, p = 0.13; **Figure 1G**). Collectively, these results indicate a DLS-specific upregulation of EAAT2 in overtrained mice.

The upregulation of EAAT2 in the DLS is consistent with the role of this striatal region in controlling habitual, inflexible behavior following task overtraining (Dayan and Balleine, 2002; Shiflett and Balleine, 2011b). As EAAT2 affects glutamatergic synaptic plasticity processes (Valtcheva and Venance, 2016), this finding raises the possibility that EAAT2 regulation influences defined aspects of action control, such as representing a permissive mechanism to enable the updating of A-O associations, which is required for efficient performance during the omission procedure. Regulation of EAAT2 expression might be less important to adapt to outcome devaluation, as this does not require the learning of novel environmental contingencies.

### Activation of Gq signaling in DLS astrocytes reduces EAAT2 expression in overtrained mice and preserves behavioral flexibility

Astrocytes express ~85% of total brain EAAT2 (Petr et al., 2015), which suggests that the upregulation of EAAT2 in overtrained mice primarily occurs in these cells. To determine whether astrocytic regulation of EAAT2 expression is specifically associated with the inability to encode changes in A-O associations that follows task overtraining, we first established a chemogenetic approach to alter EAAT2 expression in DLS astrocytes. EAAT2 internalization and degradation is controlled by the Gq signaling effector protein kinase C (PKC) (González-González et al., 2008). We reasoned that chemogenetic activation of astrocytic Gq signaling during overtraining should reduce EAAT2 expression: if behavioral inflexibility depends on increased EAAT2 expression in the DLS, such a reduction during overtraining should preserve the behavioral sensitivity to changes in action-outcome contingency.

Specific, *in vivo* chemogenetic stimulation of astrocytic Gq signaling can be achieved through adeno-associated virus (AAV)-mediated expression of the Designer Receptor Exclusively Activated by Designer Drugs (DREADD)-hM3Dq receptor under control of the glial fibrillary acidic protein (GFAP) promoter (Agulhon et al., 2013), followed by systemic administration of the cognate synthetic ligand clozapine-N-oxide (CNO) (Armbruster et al., 2007). We unilaterally infused AAV5-GFAP-hM3Dq (AAV-hM3Dq) in the DLS, and AAV5-GFAP-eGFP (AAV-eGFP) in the contralateral DLS (**Figure 2A**). To rule out non-specific behavioral effects of CNO, we used AAV-eGFP as a standard control (Gomez et al., 2017). We observed clear colocalization of the hemagglutinin (HA)-tagged hM3Dq with the astrocytic cytosolic marker ALDH1L1 (Cahoy et al., 2008), but not with the neuronal cytosolic marker MAP2 (Soltani et al., 2005), confirming specific hM3Dq expression in astrocytes (mice n = 6, ALDH1L1, 0.8 ± 0.01; MAP2, 0.18 ± 0.04; % of cells expressing hM3Dq, t-test, t = 14.54, df = 12, **** p < 0.0001; **Figure 2B** and **Figure S2A**). We then measured mRNA levels of the immediate early gene *cFos* and performed cFOS immunohistochemistry upon intraperitoneal administration (i.p.) of CNO (3 mg/kg), a strategy previously used to validate a similar chemogenetic approach (Yang et al., 2015). *In vivo* CNO administration increased *cFos* mRNA in the AAV-hM3Dq-infused hemispheres (AAV-eGFP n = 6, 100% ± 11.96, AAV-hM3Dq n = 6, 455% ± 58.90; *cFos* expression, paired t-test, t = 7.2, df = 5, *** p = 0.0008; **Figure 2C**). cFOS immunoreactivity was also greater in hM3Dq-positive cells than in NeuN-positive cells (mice n = 5, 647 cells, NeuN^+^, 10.1% ± 3.5; HA-hM3Dq^+^, 88% ± 2.3; % of cells expressing cFOS, independent samples t-test, t = 18.7, df = 8, **** p < 0.0001; **Figure 2D**), indicating that chemogenetic activation of GFAP-expressed hM3Dq predominantly elicits cFOS responses in astrocytes.

**Figure 2.**
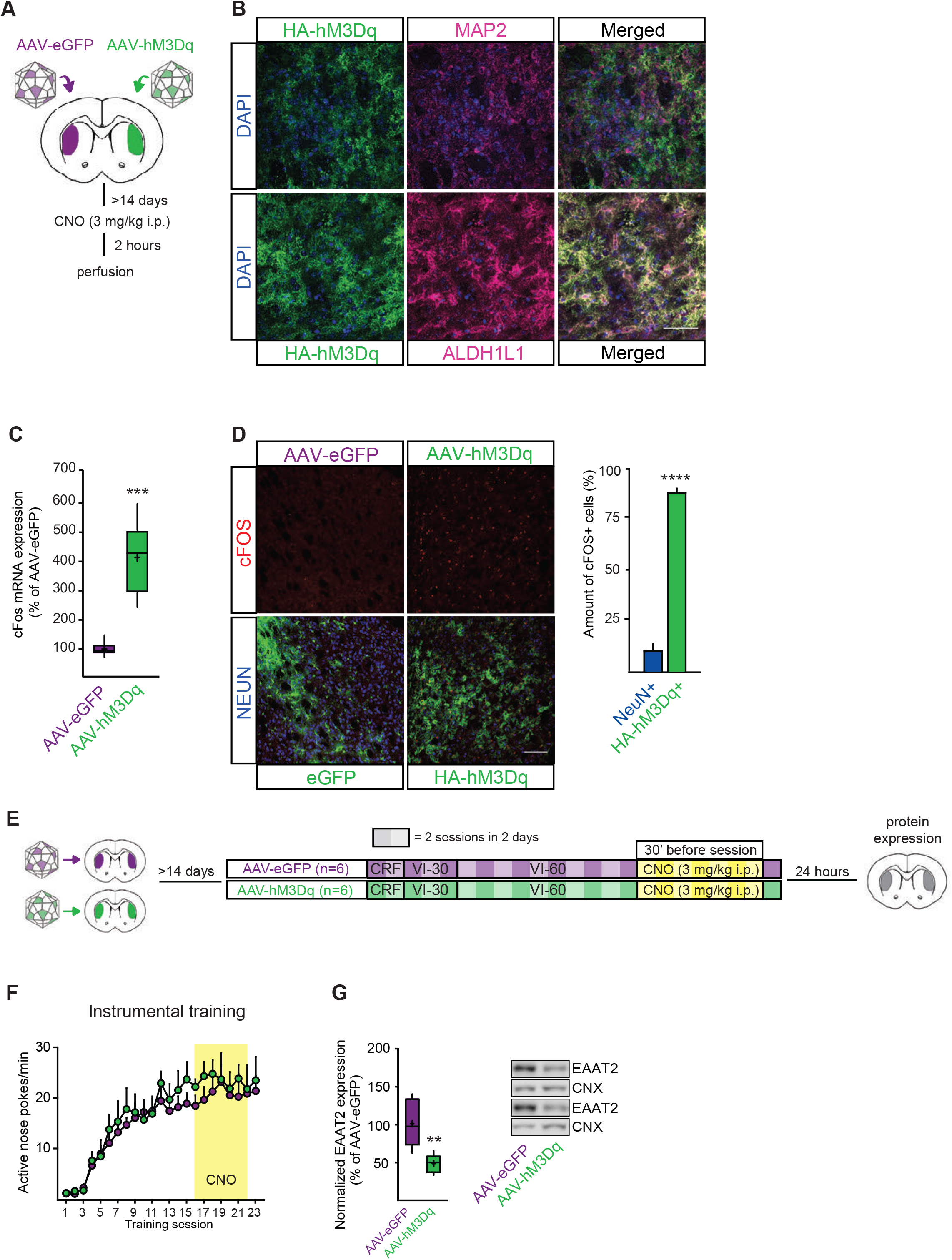
Chemogenetic activation of astrocytic hM3Dq in the DLS reduces EAAT2 expression. (**A**) Schematic of experimental design. (**B**) Representative collapsed confocal Z-stacks of immunofluorescent detection of (Left) HA-hM3Dq (AAV-GFAP-hM3Dq), (Middle) the astrocytic cytosolic marker ALDH1L1 or the neuronal cytosolic marker MAP2, and (Right) overlays. Scale bar is 50 μm. (**C**) Box-whisker plot of normalized *cFos* mRNA levels of AAV-eGFP and AAV-hM3Dq hemispheres, 30 min after clozapine-N-oxide (CNO) administration (3 mg/kg). (**D**) (Left) Representative collapsed confocal Z-stacks of immunofluorescent detection of cFOS, NeuN and HA-hM3Dq or eGFP expression 2 h after CNO administration. (Right) Quantified cFOS expression in HA-positive and NeuN-positive cells. Scale bar is 100 μm. (**E**) Schematic depicts the experimental time course. CNO was administered 30 min before the VI-60 training sessions. (**F**) ANP rates of AAV-eGFP and AAV-hM3Dq animals. (**G**) Box-whisker plot of normalized EAAT2 protein levels in the DLS of AAV-eGFP and AAV-hM3Dq mice; insets are two sample blots from each group. CNX, calnexin. (**D,F**) In the averaged time courses and bar graphs, values are presented as mean ± S.E.

To confirm that the activation of astrocytic Gq signaling during overtraining reduces EAAT2 protein expression, we bilaterally infused AAV-eGFP or AAV-hM3Dq into the DLS (**Figure 2E**). Two weeks after surgery, these mice were overtrained to nose poke for food and received CNO (3 mg/kg, i.p.) during the late phase of training (30 min before sessions 16 to 22). We have previously shown that manipulation of glutamatergic corticostriatal activity during this time window affects instrumental control (goal-directed versus habitual) (Nazzaro et al., 2012). In AAV-hM3Dq compared to AAV-eGFP control mice, CNO administration during late training altered neither ANP rates (AAV-eGFP n = 6, AAV-hM3Dq, n = 6; ANP/min, session: F_22, 220_ = 26.06, p < 0.0001; group: F_1,10_ = 0.39, p = 0.54; session X group interaction: F_22, 220_ = 0.4, p = 0.99; **Figure 2F**), nor INP or ME rates (p ˃ 0.05; **Figure S2B**), indicating that acute activation of astrocytic hM3Dq has no effect on instrumental performance per se. However, 24 h after the last training session, EAAT2 protein was significantly lower in AAV-hM3Dq mice than in AAV-eGFP mice (AAV-eGFP n = 6, 100% ± 12.6; AAV-hM3Dq n = 6, 48.3% ± 4.9; EAAT2 expression, independent samples t-test, t = 3.8, df = 10, ** p = 0.0034; **Figure 2G**). These results demonstrate that chemogenetic activation of astrocytic Gq signaling in the DLS is an effective strategy to reduce the overtraining-induced rise in EAAT2 protein levels.

We used this experimental strategy to investigate if activation of astrocytic Gq activity restores behavioral flexibility in overtrained mice. As before, we bilaterally infused mice with AAV-hM3Dq or AAV-eGFP in the DLS, and exposed them to overtraining and CNO treatment (3 mg/kg, i.p., 30 min before sessions 16 to 22. To control for any non-specific behavioral effects of hM3Dq expression in the DLS, an additional experimental group was bilaterally injected with AAV-hM3Dq, trained and treated with saline instead of CNO (**Figure 3A, E**). ANP rates during training were similar among groups (AAV-eGFP + CNO n = 6; AAV-hM3Dq + saline n = 7; AAV-hM3Dq + CNO n = 10; ANP/min, session: F_22, 440_ = 60.83, p < 0.0001; group: F_2,20_ = 0.43, p = 0.66; session X group interaction: F_44, 440_ = 1.04, p = 0.4; **Figure 3B**), as were INP and ME rates (p ˃ 0.05; **Figures S3C-D**). During the omission procedure, only AAV-hM3Dq-infused mice treated with CNO demonstrated behavioral sensitivity to a change in the action-outcome contingency (ANP/min, contingency: F_1,20_ = 2.8, p = 0.11; group: F_2,20_ = 1.05, p = 0.37; contingency X group interaction: F_2,20_ = 5.38, p = 0.014; Sidak, positive versus negative contingency: AAV-eGFP + CNO, p = 0.88; AAV-hM3Dq + saline = 1; AAV-hM3Dq + CNO, ** p = 0.002; **Figure 3C**). Time course analysis of ANP ratios reveals a main group effect between the three mouse cohorts (ANP/min, time: F_5,100_ = 0.46, p = 0.80; group: F_2,20_ = 7.88, ** p = 0.003; interaction: F_10,100_ = 1.2, p = 0.3; time X group interaction: F_10,100_ = 1.24, p = 0.27; **Figure 3C**). Accordingly, AAV-hM3Dq mice treated with CNO received a higher number of pellets than the other two groups during A-O reversal **(Figure S3B)**. This indicates more flexible behavioral control, as the number of pellets obtained during the control session did not differ among groups (contingency: F_1,20_ = 108.30, p < 0.0001; group: F_2,20_ = 7.07, p = 0.005 contingency X group interaction: F_2,20_ = 5.1, p = 0.02; Sidak, positive contingency: AAV-eGFP + CNO versus AAV-hM3Dq + saline, p > 0.9999; AAV-eGFP + CNO versus AAV-hM3Dq + CNO, p = 0.99; AAV-hM3Dq + saline versus AAV-hM3Dq + CNO, p = 0.99; Sidak, negative contingency: AAV-eGFP + CNO versus AAV-hM3Dq + saline, p = 0.91; AAV-eGFP + CNO versus AAV-hM3Dq + CNO, *** p = 0.003; AAV-hM3Dq + saline versus AAV-hM3Dq + CNO, ** p = 0.0012; **Figure S3B**). To further confirm the specificity of our chemogenetic approach, astrocytic Gq activation was further validated by GFAP immunohistochemistry 2 h after completion of the post-training omission procedure. GFAP immunofluorescence was significantly higher in AAV-hM3Dq animals treated with CNO than in control groups (fluorescent intensity, 1WA, F_2,9_ = 19.31, p = 0.0006; **Figure 3D**). Taken together, these results show that specific activation of astrocytic Gq signaling in the DLS during the late phase of overtraining reduces EAAT2 expression and restores behavioral flexibility.

**Figure 3.**
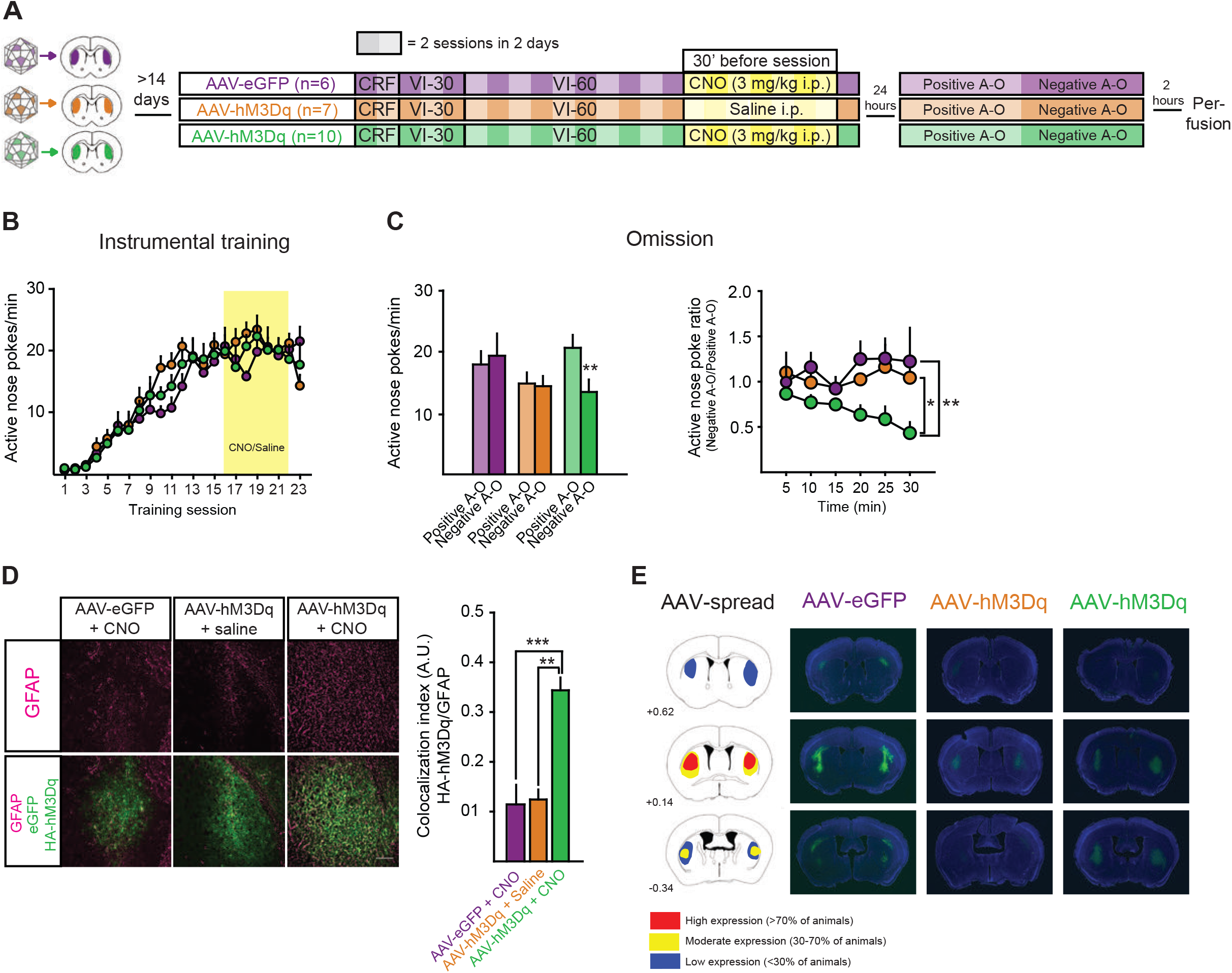
Chemogenetic activation of astrocytic hM3Dq in the DLS preserves behavioral flexibility in overtrained mice. (**A**) Schematic of behavioral and pharmacological regimens. (**B**) ANP rates during training (AAV-eGFP+CNO n = 6; AAV-hM3Dq + saline n = 7; AAV-hM3Dq + CNO n = 10. (**C**) Post-training omission procedure. (Left) Comparison of ANP rates under positive and omission contingencies (post hoc Sidaks on contingency effect: AAV-eGFP + CNO, positive contingency: 17.9 ± 2.2, negative contingency: 19.3 ± 3.6, p = 0.88; AAV-hM3Dq + saline, positive contingency: 14.8 ± 1.8, negative contingency: 14.4 ± 1.7, p = 0.99; AAV-hM3Dq + CNO, positive contingency: 22.8 ± 3, negative contingency: 16.4 ± 2.8, ** p = 0.002). (Right) Comparison of ANP ratios across experimental groups (post hoc Sidaks on main group effect: AAV-eGFP + CNO versus AAV-hM3Dq + saline, p = 0.89; AAV-hM3Dq + saline versus AAV-hM3Dq + CNO, * p = 0.02; AAV-eGFP + CNO versus AAV-hM3Dq + CNO, ** p = 0.006). (**D**) (Left) Representative collapsed confocal Z-stacks of GFAP immunoreactivity after post-training omission test. Scale bar is 200 μm. (Right) GFAP-expression was significantly increased in AAV-hM3Dq + CNO animals (1WA post hoc Tukey, AAV-eGFP + CNO versus AAV-hM3Dq + saline, p > 0.05; AAV-hM3Dq + saline versus AAV-hM3Dq + CNO, ** p < 0.01; AAV-eGFP + CNO versus AAV-hM3Dq + CNO, *** p < 0.001). (**B**-**D**) In the averaged time courses and bar graphs, values are presented as mean ± S.E. (**E**) (Left) Visualization of average spread of AAV-injections across all animals included in the analysis. (Right) AAV-spread of representative animals for each of the three experimental groups.

### Transient knockdown of EAAT2 in the DLS preserves behavioral flexibility in overtrained mice

Our data suggest that astrocytes regulate EAAT2 expression to influence instrumental control. That is, overtraining is associated with upregulation of EAAT2 and results in inflexible control of behavior (i.e. behavioral insensitivity to A-O contingency reversal), whereas activation of astrocytic Gq signaling during the late phase of training both reduces EAAT2 expression and preserves behavioral flexibility. Nevertheless, activation of astrocytic Gq signaling could prevent inflexible behavior via mechanisms other than the modulation of EAAT2 protein levels, for example, via the release of gliotransmitters (Scofield et al., 2015; Yang et al., 2015).

To test whether the regulation of EAAT2 protein levels is necessary for the expression of overtraining-induced behavioral inflexibility, we transiently reduced EAAT2 protein in the DLS during the late phase of training using Vivo-Morpholinos (MOs). MOs reduce the expression of specific proteins by preventing the formation of the ribosomal initiation complex and have been used to transiently downregulate EAAT2 levels (Reissner et al., 2015; Reissner et al., 2012). First, we confirmed that EAAT2 protein is specifically reduced by unilaterally infusing MOs targeting EAAT2 (EAAT2 MO, 3 pmol) in the DLS on three consecutive days, while the contralateral DLS received control MOs (3 pmol; **Figure 4A**) (Eisen and Smith, 2008). Seven days after the last infusion, EAAT2 protein levels were significantly lower in the DLS injected with EAAT2 MO than in the control MO-infused DLS (Control MO n = 6, 100% ± 7.1; EAAT2 MO n = 6, 60.4% ± 8.6; EAAT2 expression, Wilcoxon matched-pairs test, W = −21, * p = 0.03; **Figure 4B**). Twelve days after the last MO infusion, there was no difference between groups (Control MO n = 6, 100% ± 10.2; EAAT2 MO n = 6, 104.8% ± 10.2; EAAT2 expression, Wilcoxon matched-pairs test, W = 5, p = 0.69; **Figure 4B**). In the DMS, EAAT2 protein levels were comparable in the EAAT2 MO- and control MO-injected groups 7 days after the last MO infusion (Control MO n = 6, 100% ± 10; EAAT2 MO n = 6, 115% ± 10.9; Wilcoxon matched-pairs test, W = 9, p = 0.44; **Figure 4B**). In summary, these experiments indicate that *in vivo* MO infusion in the DLS transiently reduces EAAT2 expression in the DLS without affecting the adjacent DMS.

**Figure 4.**
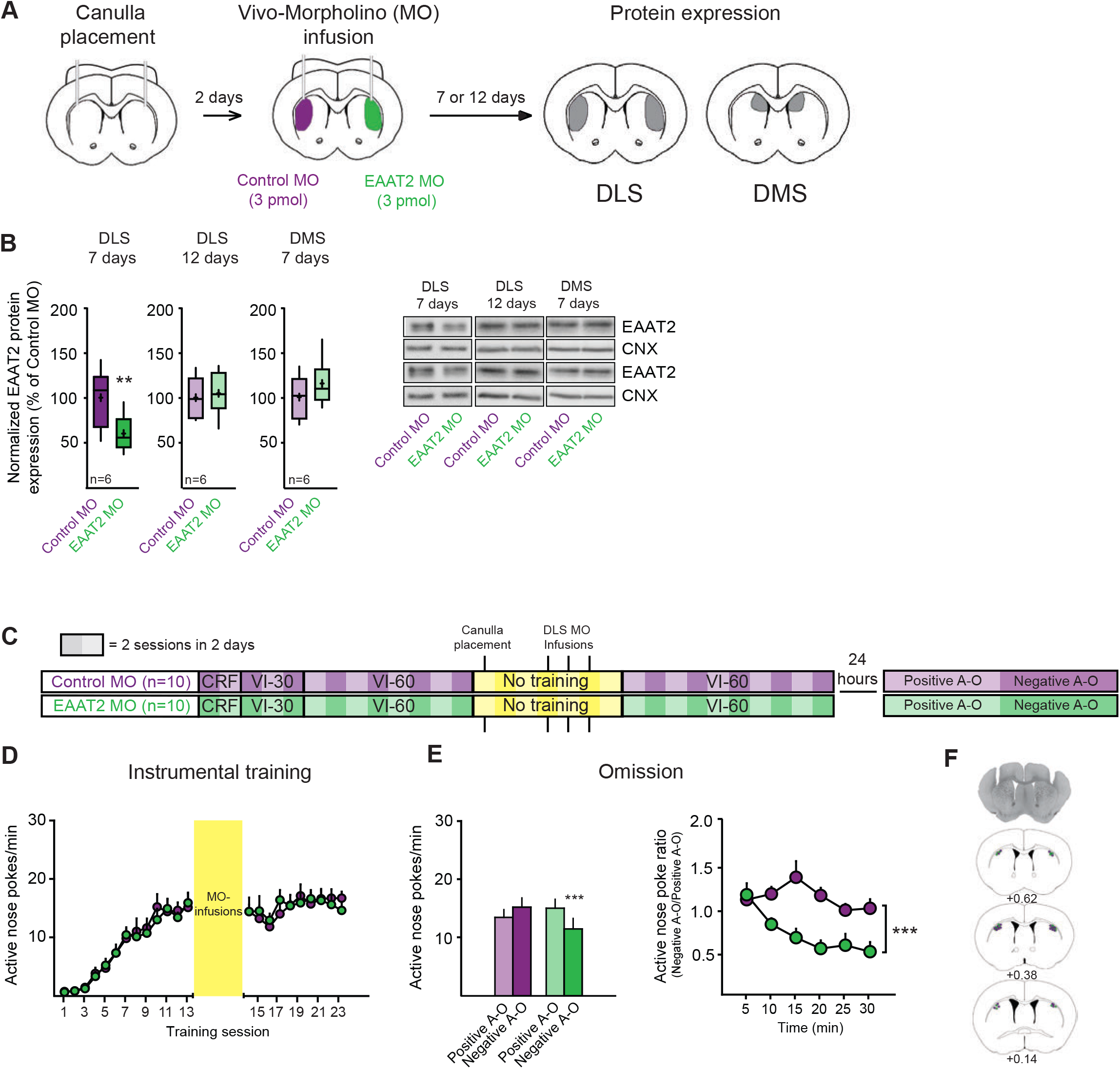
Transient knockdown of EAAT2 in the DLS impairs inflexible behavior. (**A**) Schematic of the experimental design to validate the transient knockdown of EAAT2 by Vivo-Morpholino (MO). (**B**) Normalized EAAT2 expression in the DLS (7 and 12 days) and in the DMS (7 days) after the last MO infusion. Insets are two representative EAAT2 blots from EAAT2 MO- and control MO-injected DLS. CNX, calnexin. (**C**) Schematic of behavioral regimens and DLS MO infusions. (**D**) ANP rates across training of control MO- and EAAT2 MO-infused animals. (**E**) Post-training omission procedure. (Left) Comparison of ANP rates under positive and omission contingencies for both control and EAAT2 MO groups (post hoc Sidaks on main contingency effect: Control MO n =10, positive contingency: 13.5 ± 1.3, negative contingency: 15.1 ± 1.5, p > 0.05; EAAT2 MO n =10, positive contingency: 15.2 ± 1.6, negative contingency: 11.5 ± 1.8, *** p < 0.001).

Next, we tested the effect of EAAT2 knockdown in the DLS on behavioral flexibility. We trained mice to nose poke for food for 15 sessions, after which we bilaterally implanted infusion cannulas in the DLS to allow local MO administration (**Figure 4C, F**). To knockdown EAAT2 expression during late training sessions, we administered EAAT2 MOs or control MOs for three consecutive days and resumed training two days after the last infusion. MO infusion had no effect on ANP (ANP/min; Control MO n = 10, EAAT2 MO n = 10, ANP/min, session: F_22,396_ = 40.39, p < 0.0001; group: F_1,18_ = 0.007, p = 0.93; session X group interaction: F_22,396_ = 0.41, p = 0.99; **Figure 4D**), INP or ME rates during training (**Figures S4C, D**). However, during the omission paradigm, administration of EAAT2 MO significantly affected the behavioral response. While control MO-treated mice did not reduce the number of nose pokes after the A-O contingency reversal, EAAT2 MO-treated mice did (Control MO n = 10, EAAT2 MO n = 10, ANP/min, contingency: F_1,18_ = 5.63, p = 0.03; group: F_1,18_ = 0.13, p = 0.72; contingency X group interaction: F_1,18_ = 24.33, p = 0.0001; Sidak, positive versus negative contingency: Control MO, p > 0.05; EAAT2, *** p < 0.001; **Figure 4E**). Two-way ANOVA analysis of ANP ratio time courses shows a main group effect of MO treatment (time: F_5,90_ = 6.33, p < 0.0001; group: F_1,18_ = 15.39, *** p = 0.001; time X group interaction: F_5,90_ = 4.98, p = 0.0005, **Figure 4E**). There were no differences in INP or ME rates between control and A-O reversal sessions in the two experimental groups (**Figure S4C, D**). Accordingly, EAAT2 MO mice received more reinforcers than control MO-treated mice during A-O reversal (obtained reinforcers, contingency: F_1,18_ = 23.23, p = 0.0001; group: F_1,18_ = 4.64, p = 0.05; contingency X group interaction: F_1,18_ = 3.43, p = 0.08; Sidak, Control MO versus EAAT2 MO: positive contingency, p = 0.99; negative contingency, * p = 0.02; **Figure S4B**).

To exclude non-specific behavioral effects due to general cell death upon MO infusion, we quantified astrocyte reactivity and cell nuclei through NeuN and GFAP staining in the DLS. There were no differences between control MO and EAAT2 MO groups (NeuN expression: Control MO, n = 5, 100% ± 2.5; EAAT2 MO, n = 5, 105.3% ± 2.6; Mann-Whitney Test, U = 6, p = 0.22; GFAP expression: Control MO, n = 5, 100% ± 15; EAAT2 MO, n = 5, 74.7% ± 7; Mann-Whitney Test, U = 5.5, p = 0.17; **Figure S4E**).

Collectively, these results show that during the late phase of training transient knockdown of EAAT2 in the DLS interferes with the expression of behavioral inflexibility, further supporting that experience-dependent plasticity of EAAT2 plays a key role in performance control by encoded A-O contingencies.

## Discussion

This study demonstrates that astrocytic regulation of EAAT2 expression in the DLS shapes flexible control of behavior. We find that loss of behavioral flexibility upon task overtraining in a contingent operant paradigm (i.e. nose poke for food reinforcer) is associated with upregulation of EAAT2 expression in this striatal region.

Behavioral flexibility, which characterizes goal-directed actions, involves the inhibition of ongoing behaviors, as well as the acquisition and execution of alternative behavioral strategies (Bonnavion et al., 2019). Our results show that overtraining-induced upregulation of EAAT2 expression in the DLS is directly linked to response perseverance despite a reversal of the prevailing A-O contingency. This indicates a loss of behavioral flexibility upon task overtraining, consistent with a transition from a goal-directed to a habitual action (Corbit et al., 2012; Coutureau and Killcross, 2003; Gremel et al., 2016; Gremel and Costa, 2013; Yin et al., 2004, 2006; Yin et al., 2005b). Apart from a reduced sensitivity to changes in A-O contingencies, habitual control of behavioral performance is also characterized by an inability to update changes in the value of task outcomes (Balleine and Dickinson, 1998). Indeed, the devaluation test in overtrained mice (**Figure S1C**) reveals this insensitivity to changes in the outcome value. While our experimental design assesses the impact of EAAT2 protein levels on one of the two aspects of habit expression (i.e. reduced sensitivity to novel A-O contingencies), it does not allow the conclusive demonstration of how experience-dependent regulation of astrocytic EAAT2 expression contributes to habit learning. Our omission procedure assesses the ability to erase a previously-learned A-O association (extinction of nose poking), and/or the acquisition of a novel A-O association (learning to refrain from nose poking to obtain a reinforcer), and not behavioral sensitivity to altered outcome values. Despite this limitation, our results nevertheless provide mechanistic insights into how A-O associations are maintained at frontostriatal circuits: we establish that glutamate transporter plasticity in the DLS is a cellular substrate that supports overtraining-induced behavioral inflexibility.

That task overtraining induces changes to EAAT2 expression in the DLS, but not in the DMS, indicates an altered net excitatory input to the DLS after overtraining. This finding is in line with observations that regulation of glutamatergic synapses in the DLS is crucial to maintain flexible control over behavior (O’Hare et al., 2016; Shiflett and Balleine, 2011a). The timing and magnitude of glutamate transients in the synaptic cleft is shaped by astrocytic EAAT2, a process that determines glutamate receptor activation in response to specific patterns of presynaptic activity (Bergles and Jahr, 1997; Clements et al., 1992). EAAT2 also contributes to the regulation of glutamate spillover between synapses, and thereby preserves the specificity of glutamatergic transmission and concurrent plasticity (Martin-Fernandez et al., 2017; Valtcheva and Venance, 2016). It has previously been shown that plasticity at glutamatergic cortical inputs to the DLS is associated with loss of behavioral flexibility (Gremel and Costa, 2013; Nazzaro et al., 2012; O’Hare et al., 2016). In addition, timing-dependent LTD (t-LTD) at corticostriatal synapses on striatal projection neurons (SPNs) is impaired by pharmacological upregulation of EAAT2 expression (Valtcheva and Venance, 2016). Increased striatal EAAT2 expression is expected to enhance glutamate uptake and reduce glutamate spillover, likely influencing the activity of group I metabotropic glutamate receptors 1/5 (mGluR1/5) and downstream signaling pathways, including endocannabinoid (eCB) biosynthesis (Heifets and Castillo, 2009). The loss of eCB-mediated LTD in DLS SPNs is directly associated with behavioral inflexibility induced by chronic exposure to Δ-9-THC or overtraining of nose poking for food (Nazzaro et al., 2012). While the current study does not directly address the relationship between glutamate uptake and behavioral inflexibility, our collective results suggest that training-induced upregulation of EAAT2 in the DLS may negatively interfere with the ability to encode changes in A-O association by affecting mGluR1/5-eCB-dependent regulation of synaptic plasticity.

Notably, the ensemble activity pattern of neurons steadily increases in the DLS as training progresses (Thorn et al., 2010; Yin et al., 2009), which might be supported by the loss of processes regulating synaptic mechanisms of depression at corticostriatal inputs. In addition to eCB-mediated LTD, this pattern could also involve the regulation of mGluR2/3 receptors expressed on cortical terminals, whose activation decreases the release probability of glutamate (Niswender and Conn, 2010). In this model, increased glutamate uptake through EAAT2 in overtrained mice might result in reduced activation of mGluR2/3 receptors and increased strength of cortical inputs to the DLS, thereby affecting the entrainment of task-related SPNs there.

Regulation of instrumental performance (i.e. habitual control of behavior) by ambient glutamate has been previously associated with psychostimulant administration (Corbit et al., 2014). Although EAAT2 expression levels were not directly measured, the study provides evidence for changes in striatal synaptic plasticity, thus suggesting that EAAT2 is involved in drug-induced behavioral alterations (Kalivas, 2009). Furthermore, induced deficiency of astrocytic EAAT2 in the striatum has been associated with pathological repetitive behaviors (Aida et al., 2015). Here, we show that increased EAAT2 expression is directly linked to action inflexibility induced by task overtraining, under physiological conditions. Results obtained in EAAT2 knockout mice and in mice exposed to an instrumental learning paradigm (this study) may not be directly comparable, as the genetic deletion of astrocytic EAAT2, even conditionally, might induce adaptive cellular mechanisms. Moreover, in the knockout model, astrocytic EAAT2 deficiency was induced brain-wide and not specifically in the DLS. We can also speculate that opposite changes in EAAT2 protein levels, likely resulting in either increased (EAAT2 downregulation) or decreased (EAAT2 upregulation) glutamate spillover, might affect corticostriatal inputs by targeting different forms of plasticity. That is, EAAT2 inhibition might lead to an aberrant form of LTP at corticostriatal synapses, as recently shown (Valtcheva and Venance, 2016). In contrast, EAAT2 overexpression might lead to a loss of LTD by primarily affecting mGluR1/5-mediated signaling or may cause enhanced glutamate release by reducing activation of mGluR2/3R. Thus, either aberrant LTP or a loss of LTD may compromise the dynamic range of synaptic activity in the DLS and promote inflexible behavioral strategies. These possibilities nevertheless support the notion that fine-tuned regulation of EAAT2 expression is crucial for establishing appropriate glutamate dynamics at striatal circuits.

Our results show that chronic stimulation of Gq-coupled downstream G-protein coupled receptors (GPCR) normalizes EAAT2 expression in overtrained mice. GPCR regulation of glutamate uptake coincides with the structural reorganization of astrocytic processes, raising the possibility that astrocytes integrate synaptic information to spatially regulate EAAT2 expression (Bernardinelli et al., 2014; Genoud et al., 2006; Sweeney et al., 2017). Plasticity in the astrocytic coverage of synapses facilitates the response to environmental stimuli and suggests that activity-dependent remodeling in the structure of astrocytic processes underlies behavioral adaptations (Oliet et al., 2001; Piet et al., 2004). The present work does not provide direct evidence that structural plasticity in astrocytic processes is associated with habitual behavior. However, since structural plasticity in astrocytes is regulated by Gq-signaling (Bernardinelli et al., 2014), coincides with increased EAAT2 expression (Genoud et al., 2006), and facilitates the response to environmental stimuli (Piet et al., 2004), an intriguing possibility is that astrocytes might regulate spatial expression patterns of EAAT2 to exert local control over glutamate dynamics and accommodate the synaptic changes that facilitate behavioral inflexibility.

In conclusion, our results establish a direct link between astrocytic regulation of EAAT2 expression and behavioral flexibility. By showing how activity in striatal astrocytes influences action control, we corroborate the importance of astrocytes in striatal functioning and the regulation of behavior.

## Funding and disclosure

This study was supported by the Fondazione Istituto Italiano di Tecnologia. The authors declare no competing financial interests.

## Author Contributions

**A.B**. Designed, performed and analyzed experiments. **R.T**. Supervised the project and directed the experiments. **A.B**. and **R.T**. wrote the manuscript.

