## Supplemental Material for "Plasticity of the glutamate transporter EAAT2 on striatal astrocytes regulates flexibility in behavior"

Figure S1

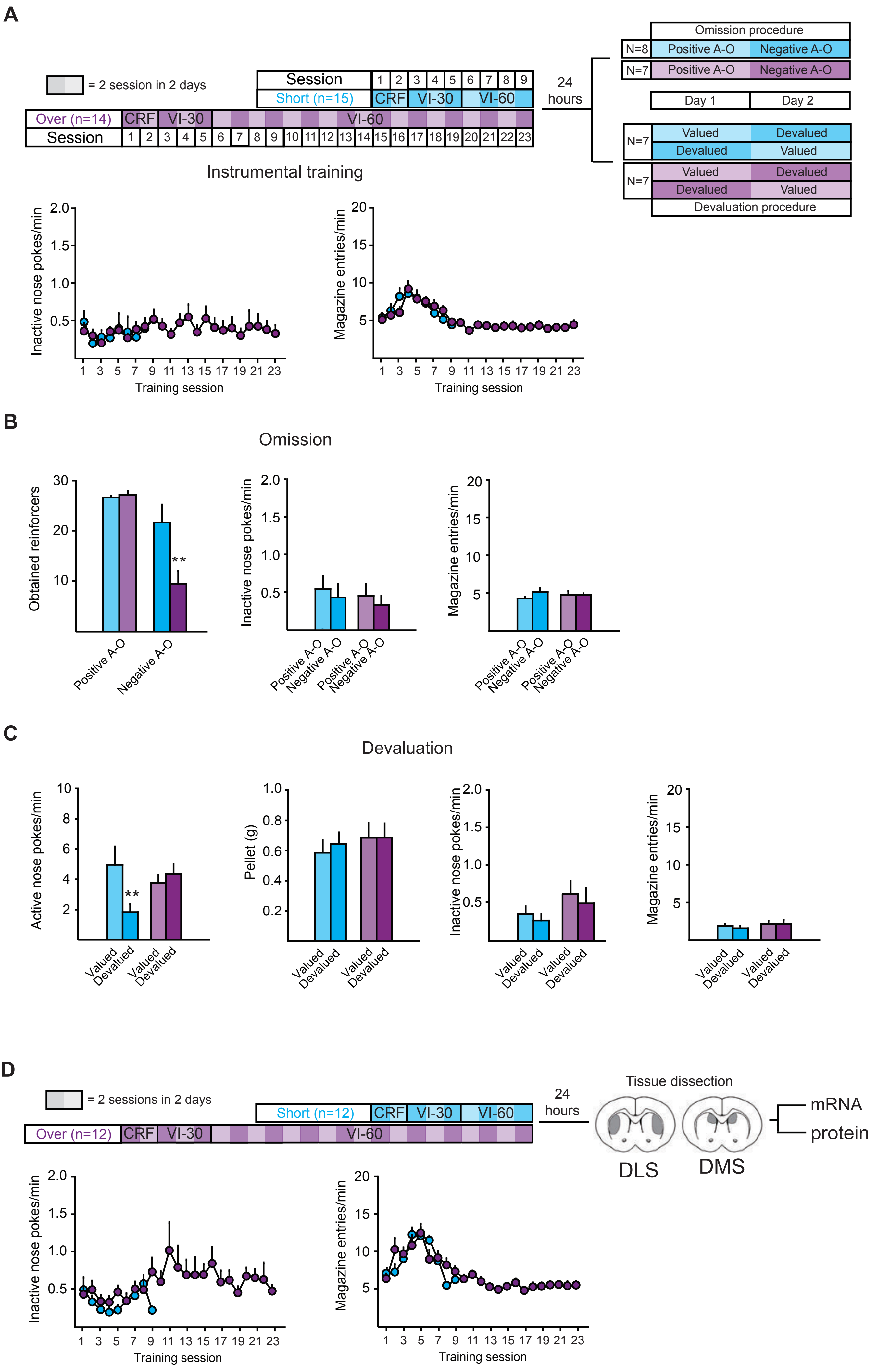

A
= 2 session in 2 days
Session
Short (n=15)
CRF
VI-30
VI-60
Over (n=14)
CRF
VI-30
VI-60
Session
24 hours
Omission procedure
N=8
Positive A-O
Negative A-O
N=7
Positive A-O
Negative A-O
Day 1
Day 2
N=7
Valued
Devalued
Devalued
Valued
N=7
Valued
Devalued
Devalued
Valued
Devaluation procedure
Instrumental training
Inactive nose pokes/min
Training session
Magazine entries/min
Training session
B
Omission
Obtained reinforcers
**
Positive A-O
Negative A-O
Inactive nose pokes/min
Positive A-O
Negative A-O
Positive A-O
Negative A-O
Magazine entries/min
Positive A-O
Negative A-O
Positive A-O
Negative A-O
C
Devaluation
Active nose pokes/min
**
Valued
Devalued
Valued
Devalued
Pellet (g)
Valued
Devalued
Valued
Devalued
Inactive nose pokes/min
Valued
Devalued
Valued
Devalued
Magazine entries/min
Valued
Devalued
Valued
Devalued
D
= 2 sessions in 2 days
Short (n=12)
CRF
VI-30
VI-60
Over (n=12)
CRF
VI-30
VI-60
24 hours
Tissue dissection
DLS
DMS
mRNA
protein
Inactive nose pokes/min
Training session
Magazine entries/min
Training session

**Figure S1**

(**A**) (Top) Schematics depict the behavioral paradigms. (Bottom) Averaged time courses (mean ± S.E.) of inactive nose poke (INP) rates (left) and magazine entry (ME) rates (right) during instrumental training (Short n = 15, Over n = 14; INP/min, session: $F_{8,216} = 1.55$, $p = 0.14$; group: $F_{1,27} = 0.001$, $p = 0.97$; session X group interaction: $F_{8,216} = 0.4$, $p = 0.92$; ME/min, session: $F_{8,216} = 6.36$, $p < 0.0001$; group: $F_{1,27} = 0.26$, $p = 0.61$; session X group interaction: $F_{8,216} = 1.58$, $p = 0.13$). (**B**) Post-training omission procedure in short- and overtrained mice (Short n = 8, Over n = 7; mean ± S.E). (Left) Amount of obtained pellets under positive and negative contingency (positive contingency: short-trained, 26.6 ± 0.4, overtrained, 27.1 ± 0.4; Sidak, $p = 0.4$; negative contingency: short-trained, 21.6 ± 3.6, overtrained, 9.4 ± 2.5; Sidak, ** $p < 0.02$). (Middle) INP rates (INP/Min, contingency: $F_{1,13} = 2.72$, $p = 0.12$; group: $F_{1,13} = 0.18$, $p = 0.68$; contingency X group interaction: $F_{1,13} = 0.01$, $p = 0.94$; short-trained: positive contingency, 0.53 ± 0.17, negative contingency, 0.42 ± 0.18; overtrained: positive contingency, 0.44 ± 0.15, negative contingency, 0.32 ± 0.11). (Right) ME rates (ME/min, contingency: $F_{1,13} = 1.39$, $p = 0.26$; group: $F_{1,13} = 0.02$, $p = 0.91$; contingency X group interaction: $F_{1,13} = 1.66$, $p = 0.22$; short-trained: positive contingency, 4.3 ± 0.3, negative contingency, 5.1 ± 0.6; overtrained: positive contingency, 4.8 ± 0.5, negative contingency, 4.7 ± 0.2). (**C**) Post-training devaluation procedure in short- and overtrained mice (Short n = 7, Over n = 7; mean ± S.E). (Left) ANP rates in the valued and devalued conditions (Short, valued condition: 5.0 ± 1.2; devalued condition 1.8 ± 0.5, Sidak: ** $p = 0.006$; Over, valued condition: 3.8 ± 0.6, devalued condition: 4.4 ± 0.7, Sidak: $p = 0.74$). (Middle, left) Pellet consumption (consumed pellets in g, condition: $F_{1,12} = 0.1$, $p = 0.76$; group: $F_{1,12} = 0.47$, $p = 0.51$; condition X group interaction: $F_{1,12} = 0.34$, $p = 0.57$; Short: valued, 0.6 ± 0.1, devalued, 0.6 ± 0.1; Over: valued, 0.7 ± 0.1, devalued, 0.7 ± 0.1). (Middle, right) INP rates (INP/min, condition: $F_{1,12} = 0.76$, $p = 0.4$; group: $F_{1,12} = 1.85$, $p = 0.2$; condition X group interaction: $F_{1,12} = 0.02$, $p = 0.9$; Short: valued, 0.4 ± 0.1, devalued, 0.3 ± 0.1; Over: valued, 0.6 ± 0.2, devalued, 0.5 ± 0.2. (Right) ME rates (ME/min, condition: $F_{1,12} = 0.25$, $p = 0.63$; group: $F_{1,12} = 0.74$, p = 0.41; condition X group interaction: $F_{1,12} = 0.38$, p = 0.55; Short, valued, 1.9 ± 0.4, devalued, 1.6 ± 0.3; Over, valued, 2.2 ± 0.4, devalued, 2.2 ± 0.5). (**D**) (Top) Schematics of training regimes followed by mRNA and protein analysis of EAAT2 expression in the DLS and DMS. (Bottom) Averaged time courses (mean ± S.E.) of INP (left) and ME rates (right) in short- (n = 12) and overtrained mice (n = 12) (INP/min, session: $F_{8,176} = 1.99$, p = 0.05; group: $F_{1,22} = 2.36$, p = 0.14; session X group interaction: $F_{8,176} = 1.77$, p = 0.09; ME/min, session: $F_{8,176} = 16.23$, $p < 0.0001$; group: $F_{1,22} = 0.18$, p = 0.68; session X group interaction: $F_{8,176} = 3.12$, p = 0.003).

### Figure S2

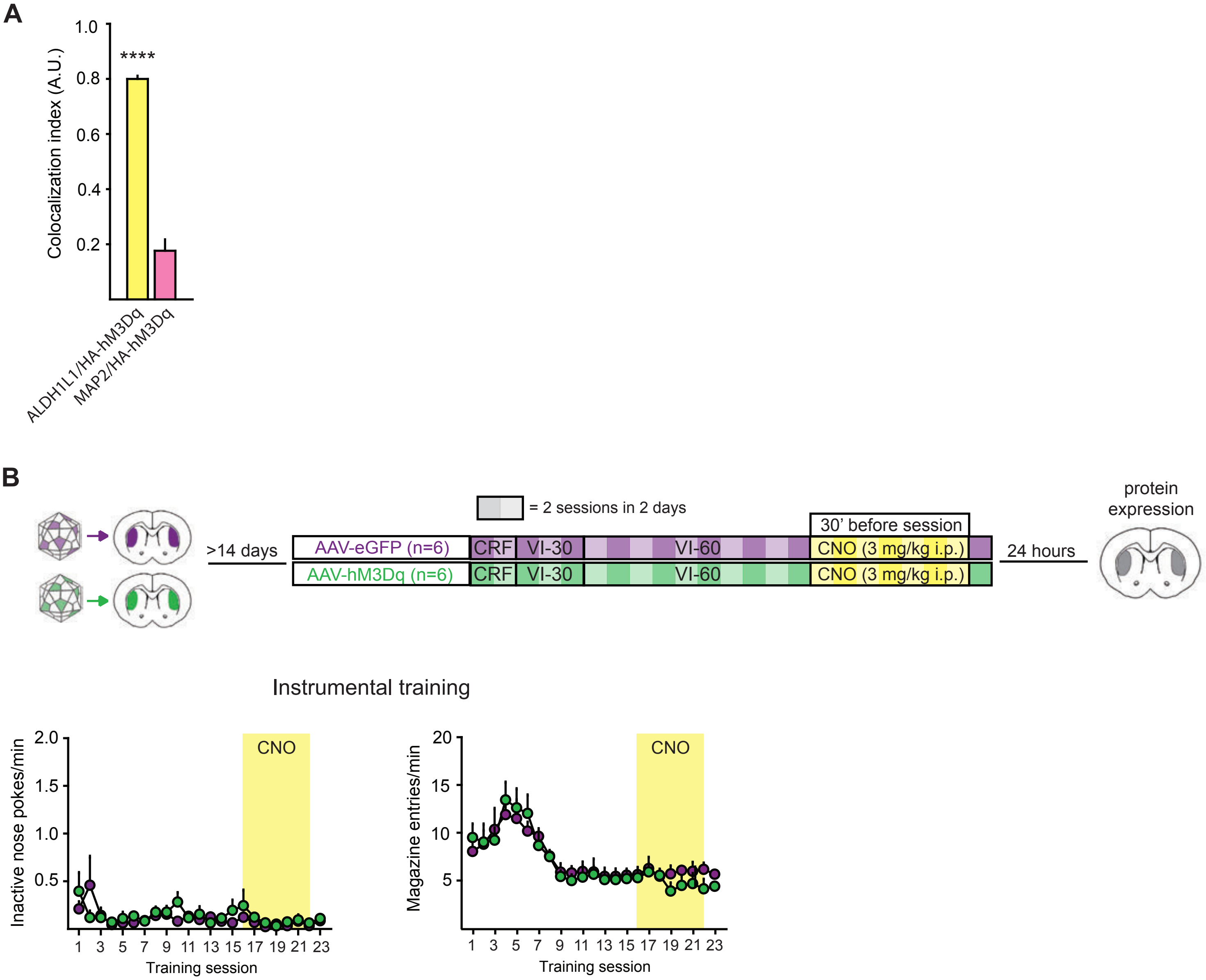

A
1.0
0.8
0.6
0.4
0.2
Colocalization index (A.U.)
****
ALDH1L1/HA-hM3Dq
MAP2/HA-hM3Dq
B
= 2 sessions in 2 days
30' before session
protein expression
>14 days
AAV-eGFP (n=6)
AAV-hM3Dq (n=6)
CRF
VI-30
VI-60
CNO (3 mg/kg i.p.)
24 hours
Instrumental training
Inactive nose pokes/min
2.0
1.5
1.0
0.5
CNO
1 3 5 7 9 11 13 15 17 19 21 23
Training session
Magazine entries/min
20
15
10
5
CNO
1 3 5 7 9 11 13 15 17 19 21 23
Training session

**Figure S2**

(**A**) Quantification of colocalization of HA-hM3Dq with ALDH1L1 or MAP2 immunofluorescence (mice n = 6). Values are expressed as mean ± S.E. **(B)** (Top) Schematics depict the behavioral paradigm and pharmacological treatments. (Bottom) Averaged time courses (mean ± S.E.) of INP rates (left) and ME rates (right) in overtrained mice injected with either the AAV-hM3Dq or AAV-eGFP viruses (AAV-hM3Dq n = 6, AAV-eGFP n = 6; INP/min, session: $F_{22,220} = 2.07$, $p = 0.005$; group: $F_{1,10} = 0.38$, $p = 0.55$; session X group interaction: $F_{22,220} = 1.131$, $p = 0.32$; ME/min, session: $F_{22,220} = 16.58$, $p < 0.0001$; group: $F_{1,10} = 0.09$, $p = 0.77$; session X group interaction: $F_{22,220} = 0.74$, $p = 0.79$).

Figure S3

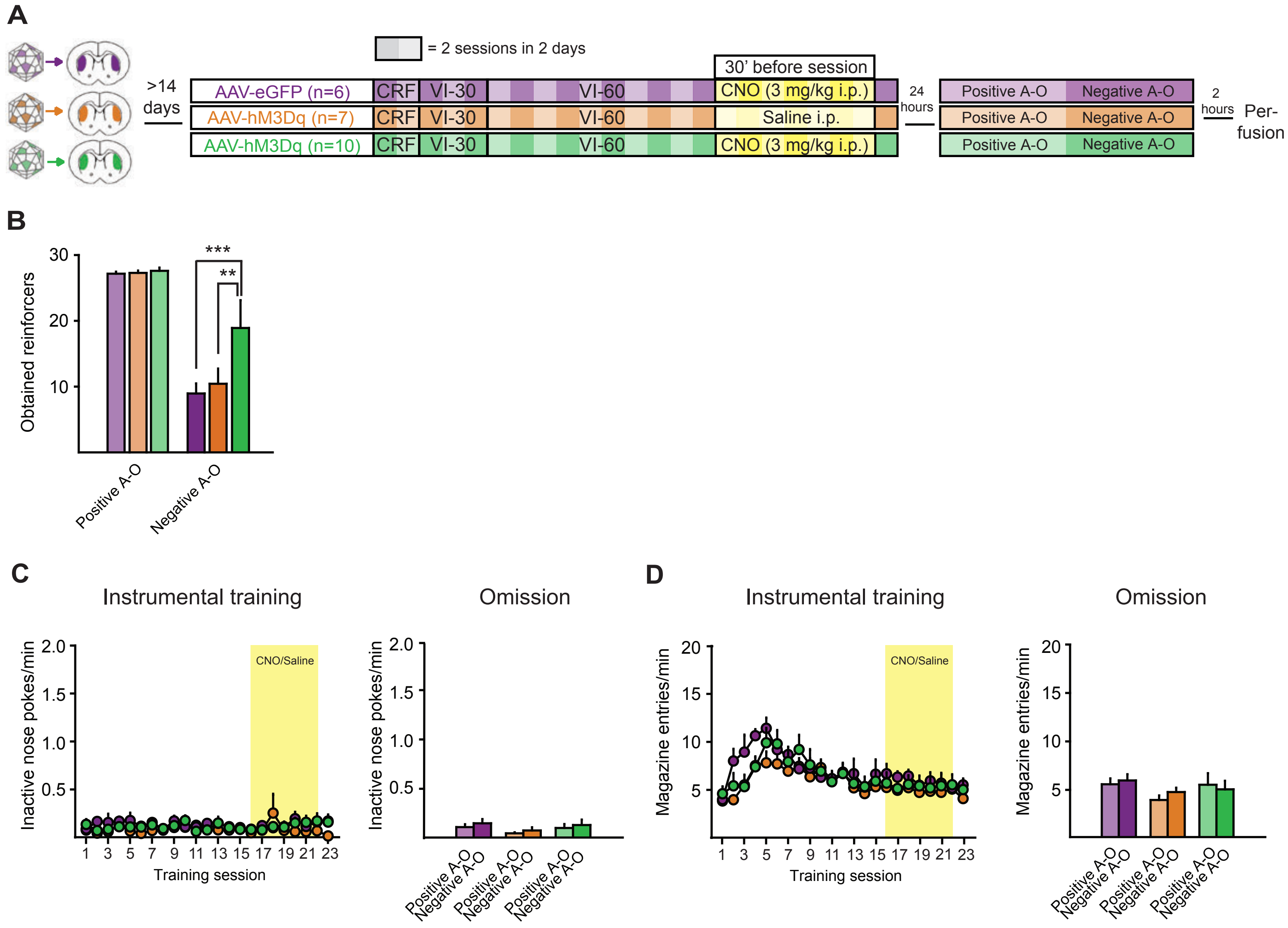

A
= 2 sessions in 2 days
>14 days
30' before session
AAV-eGFP (n=6)
AAV-hM3Dq (n=7)
AAV-hM3Dq (n=10)
CRF
VI-30
VI-60
CNO (3 mg/kg i.p.)
Saline i.p.
24 hours
Positive A-O
Negative A-O
2 hours
Per-fusion
B
Obtained reinforcers
***
**
Positive A-O
Negative A-O
C
Instrumental training
Omission
Inactive nose pokes/min
CNO/Saline
Training session
D
Magazine entries/min

**Figure S3**

(**A**) Schematics depict the behavioral paradigm and pharmacological treatments. (**B**) Number of earned pellets during the omission procedure in the three experimental groups (AAV-eGFP + CNO n = 6; AAV-hM3Dq + saline n = 7; AAV-hM3Dq + CNO n = 10 ; mean ± S.E; positive contingency: AAV-eGFP + CNO, 27.2 ± 0.3, AAV-hM3Dq + saline, 27.3 ± 0.4, AAV-hM3Dq + CNO, 27.6 ± 0.5; Sidaks: AAV-eGFP + CNO versus AAV-hM3Dq + saline, $p > 0.99$; AAV-eGFP + CNO versus AAV-hM3Dq + CNO, $p > 0.99$; AAV-hM3Dq + saline versus AAV-hM3Dq + CNO, $p > 0.99$; negative contingency: AAV-eGFP + CNO, 9.0 ± 1.5, AAV-hM3Dq + saline, 10.4 ± 2.3, AAV-hM3Dq + CNO, 18.9 ± 2.3; Sidaks: AAV-eGFP + CNO versus AAV-hM3Dq + saline, $p = 0.92$; AAV-eGFP + CNO versus AAV-hM3Dq + CNO, *** $p = 0.0003$; AAV-hM3Dq + saline versus AAV-hM3Dq + CNO, ** $p = 0.001$). (**C**) (Left) Averaged time courses (mean ± S.E.) of INP rates during instrumental training (INP/min, session: $F_{22,440} = 0.74$, $p = 0.8$; group: $F_{2,20} = 1.28$, $p = 0.3$; session X group interaction: $F_{44,440} = 0.79$, $p = 0.83$). (Right) Bar graph (mean ± S.E.) of INP rates (INP/min, contingency: $F_{1,20} = 2.53$, $p = 0.13$; group: $F_{2,20} = 0.81$, $p = 0.46$; contingency X group interaction: $F_{2,20} = 0.03$, $p = 0.97$; AAV-eGFP + CNO, positive contingency, 0.11 ± 0.03, negative contingency, 0.14 ± 0.04; AAV-hM3Dq + saline, positive contingency, 0.04 ± 0.01, negative contingency, 0.07 ± 0.03; AAV-hM3Dq + CNO, positive contingency, 0.1 ± 0.04, negative contingency, 0.13 ± 0.05). (**D**) (Left) Averaged time course ME rates (mean ± S.E.) (ME/min, session: $F_{22,440} = 9.83$, $p < 0.0001$; group: $F_{2,20} = 0.43$, $p = 0.66$; session X group interaction: $F_{44,440} = 1.14$, $p = 0.26$). (Right) Bar graph (mean ± S.E.) of ME rates (ME/min, contingency: $F_{1,20} = 2.9$, $p = 0.1$; group: $F_{2,20} = 0.68$, $p = 0.52$; contingency X group interaction: $F_{2,20} = 0.95$, $p = 0.41$; AAV-eGFP + CNO, positive contingency, 5.4 ± 0.6, negative contingency, 5.8 ± 0.6; AAV-hM3Dq + saline, positive contingency, 3.8 ± 0.5, negative contingency, 4.6 ± 0.4; AAV-hM3Dq + CNO, positive contingency, 5.3 ± 1.2, negative contingency, 5.4 ± 0.9).

### Figure S4

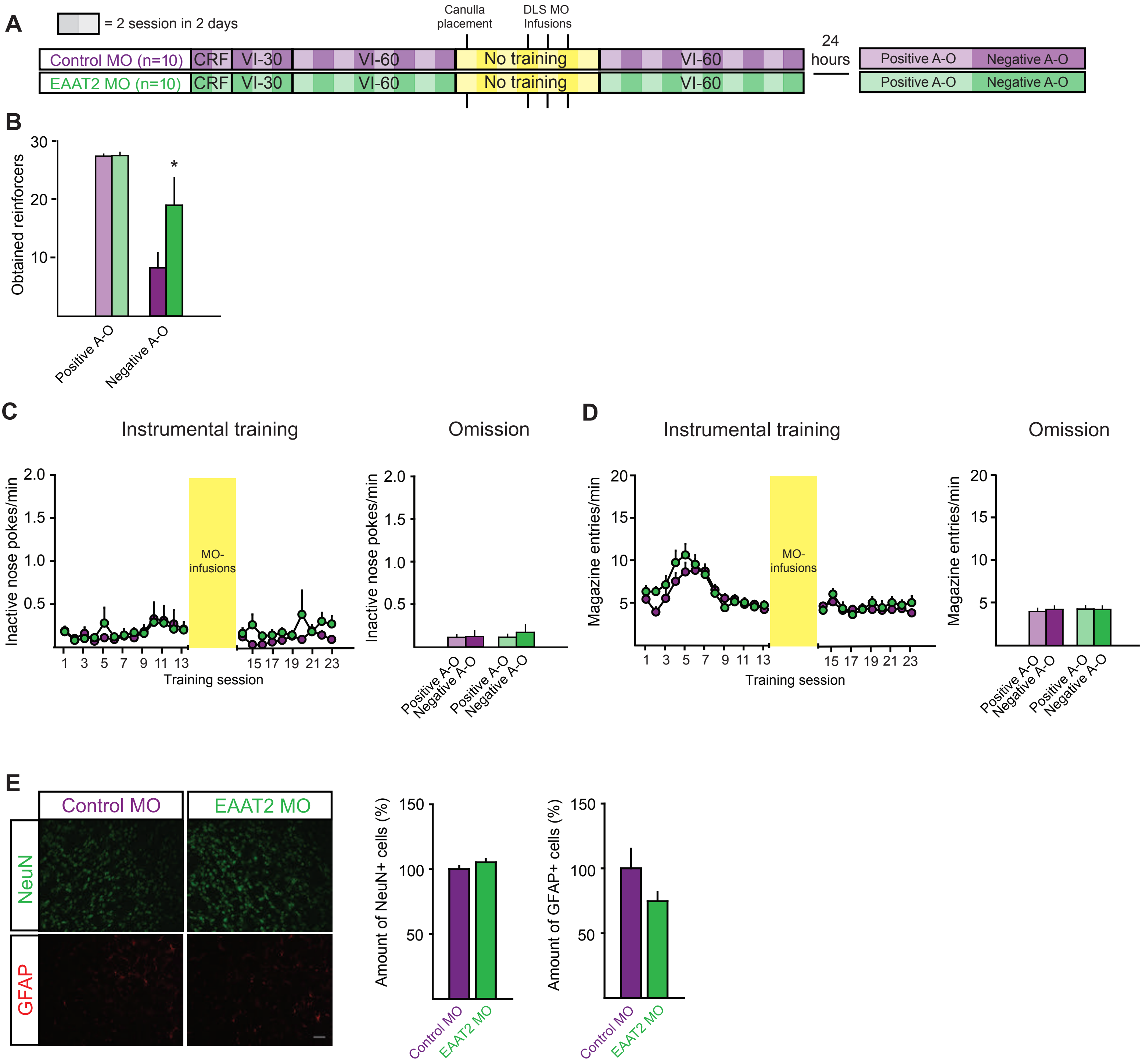

A
= 2 session in 2 days
Canulla placement
DLS MO Infusions
Control MO (n=10)
EAAT2 MO (n=10)
CRF
VI-30
VI-60
No training
24 hours
Positive A-O
Negative A-O
B
Obtained reinforcers
30
20
10
*
Positive A-O
Negative A-O
C
Instrumental training
Omission
Inactive nose pokes/min
2.0
1.5
1.0
0.5
MO-infusions
1 3 5 7 9 11 13 15 17 19 21 23
Training session
D
Magazine entries/min
20
15
10
5
E
Control MO
EAAT2 MO
NeuN
GFAP
Amount of NeuN+ cells (%)
Amount of GFAP+ cells (%)
150
100
50

**Figure S4**

(**A**) Schematics depict the behavioral paradigm and Vivo-MO infusion regimen. (**B**) Amount of obtained pellets under positive and negative contingency (positive contingency: Control MO, 27.3 ± 0.3, EAAT2 MO, 27.5 ± 0.5 Sidak, $p > 0.99$; negative contingency: Control MO, 8.5 ± 2.6; EAAT2 MO, 19.0 ± 4.7, Sidak, * $p = 0.02$). (**C**) (Left) Averaged time courses (mean ± S.E.) of INP rates (Control MO n = 10, EAAT2 MO n = 10; INP/min, session: $F_{22,396} = 1.4$, $p = 0.09$; group: $F_{1,18} = 0.49$, $p = 0.49$; session X group interaction: $F_{22,396} = 0.84$, $p = 0.68$). (Right) Bar graphs (mean ± S.E.) of INP rates (INP/min, contingency: $F_{1,18} = 1.52$, $p = 0.23$; group: $F_{1,18} = 0.15$, $p = 0.7$; contingency X group interaction: $F_{1,18} = 0.13$, $p = 0.27$, Control MO: positive contingency, 0.11 ± 0.03, negative contingency, 0.12 ± 0.03; EAAT2 MO: positive contingency, 0.12 ± 0.07, negative contingency, 0.17 ± 0.09). (**C-D**) Post-training omission procedure in Control MO and EAAT2 MO mice (Control MO n = 10; EAAT2 MO n = 10). (**D**) (Left) Averaged time courses (mean ± S.E.) of ME rates during instrumental training (ME/min, session: $F_{22,396} = 15.92$, $p < 0.0001$; group: $F_{1,18} = 0.57$, $p = 0.48$; session X group interaction: $F_{22,396} = 1.12$, $p = 0.32$). (Right) Bar graphs (mean ± S.E.) of ME rates (Control MO: positive contingency, 4 ± 0.4, negative contingency, 4.3 ± 0.4; EAAT2 MO: positive contingency, 4.3 ± 0.4, negative contingency, 4.3 ± 0.4; ME/min, contingency: $F_{1,18} = 1.33$, $p = 0.26$; group: $F_{1,18} = 0.08$, $p = 0.79$; contingency X group interaction: $F_{1,18} = 1.33$, $p = 0.26$. (**E**) Representative pictures of NeuN and GFAP staining in the DLS of Control MO- and EAAT2 MO-infused mice and relative quantification. Values are expressed as mean ± S.E.
